# Validation of the linear regression method to evaluate population accuracy and bias of predictions for non-linear models

**DOI:** 10.1101/2022.10.02.510518

**Authors:** Haipeng Yu, Rohan L Fernando, Jack CM Dekkers

## Abstract

**Background:** The linear regression method (LR) was proposed to estimate population bias and accuracy of predictions, while addressing the limitations of commonly used cross-validation methods. The validity and behavior of the LR method have been provided and studied for linear model predictions but not for non-linear models. The objectives of this study were to 1) provide a mathematical proof for the validity of the LR method when predictions are based on conditional mean, 2) explore the behavior of the LR method in estimating bias and accuracy of predictions when the model fitted is different from the true model, and 3) provide guidelines on how to appropriately partition the data into training and validation such that the LR method can identify presence of bias and accuracy in predictions.

**Results:** We present a mathematical proof for the validity of the LR method to estimate bias and accuracy of predictions based on the conditional mean, including for non-linear models. Using simulated data, we show that the LR method can accurately detect bias and estimate accuracy of predictions when an incorrect model is fitted when the data is partitioned such that the values of relevant predictor variables differ in the training and validation sets. But the LR method fails when the data are not partitioned in that manner.

**Conclusions:** The LR method was proven to be a valid method to evaluate the population bias and accuracy of predictions based on the conditional mean, regardless of whether it is a linear or non-linear function of the data. The ability of the LR method to detect bias and estimate accuracy of predictions when the model fitted is incorrect depends on how the data are partitioned. To appropriately test the predictive ability of a model using the LR method, the values of the relevant predictor variables need to be different between the training and validation sets.

## Background

Advances in high-throughput genotyping have enabled the implementation of genomic prediction, which has facilitated the genetic improvement of animals and plants based on more accurate estimated breeding values (EBV) at an early stage [e.g., 1–6]. Various genomic prediction models have been proposed and prediction performance across or within models is usually evaluated by cross-validation (CV) methods [1, 7–9]. With CV, the data set is partitioned into training and validation sets, with the training set used to fit a prediction model and estimate the breeding values of individuals in the validation set. Prediction performance is commonly evaluated with the statistic of predictivity, which is the correlation coefficient between the EBV and phenotypes adjusted for fixed effects of individuals in the validation set. Scaling predictivity by the square root of heritability (*h*^2^) provides an estimator for prediction accuracy of the EBV [10], defined as the correlation between true and estimated breeding values. While accuracy estimated with CV has been widely used to quantify the performance of genomic prediction models, pre-correcting phenotypes in the validation set using estimates of fixed effects obtained using the whole data set will overestimate the accuracy when multiple levels of fixed effects are present [11]. Additional limitations include that it can not be applied to complex models (e.g., random regression models), indirect traits (e.g., unobserved latent traits), and traits with low heritability (*h*^2^) [11].

To address these limitations of the CV methodology, Legarra and Reverter [11] proposed a linear regression (LR) method to estimate the accuracy of genomic prediction. The LR method quantifies the population accuracy and bias of predictions based on the comparison of EBV of individuals in the validation set estimated using the training data set with the EBV of those same individuals estimated using the combined training and validation sets. In the LR method literature, the training set is referred to as the partial data set (*p*) and the combined training and validation data set is referred to as the whole data set (*w*). The LR method was mathematically proven to provide unbiased estimates of the accuracy and bias of predictions for best linear unbiased prediction (BLUP) by Legarra and Reverter [11] based on results from Reverter et al. [12]. Macedo et al. [13] investigated the behavior and properties of the LR method by analyzing simulated data with pedigree-based genetic models. They studied the LR estimators of population bias and accuracy of predictions by using wrong values of *h*^2^ in the analysis and by fitting wrong models, and claimed that “the LR method works reasonably well for detection of bias when the model used is robust or close to the true model, and that it works well for estimation of accuracy even when the model is not good”. The validity and performance of the LR method for a non-linear model was explored by Bermann et al. [14]. In their study, they evaluated the performance of the LR method by fitting a threshold model to both simulation and real data sets and concluded the LR method can be useful to estimate the directions of bias, dispersion, and accuracy, though with different magnitudes. The original proof of the LR method [11] was based on the setting where the whole data set had additional phenotype records relative to the partial data set. Belay et al. [15] have recently shown that the LR method can also be applied to the setting where the whole data set has additional genotypes (rather than phenotypes) relative to the partial data set. They used the LR method to evaluate the bias and accuracy in single-step genomic predictions.

While the validity and performance of LR method has been explored using linear and non-linear models in previous studies [13, 14], a mathematical proof of its validity for non-linear methods of prediction has not yet been presented. In addition, studies about the performance of the LR method when a model other than the true model is fitted are still relatively scarce in the literature. The objectives of this study are to 1) present a mathematical proof of the validity of the LR method when predictions are based on conditional mean, regardless of whether it is a linear or non-linear function of the data 2) investigate the ability of the LR method to estimate the bias and accuracy of predictions when the fitted model differs from that used to generate the data, and 3) provide some guidelines on how to partition the data set such that the LR method can detect bias and estimate accuracy of predictions when the incorrect model is fitted.

## Theory

### Proof that 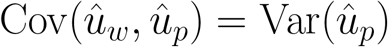

In the LR method, Legarra and Reverter [11] used var(x) to denote the variance of a random element, *x*, sampled from a single realization of the random vector (x). Here we will denote this variance by Var(*x*) = var(x). Let *u* denote the breeding value of a validation animal, and 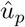 and 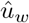 denote the estimated breeding value of u obtained from partial data and whole data, respectively. Legarra and Reverter [11] proposed the LR method for BLUP by showing 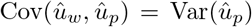 using the results from Reverter et al. [12] and assumptions of 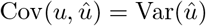 and 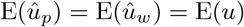. In the following, we prove the validity of the LR method for non-linear models by generalizing the proof of 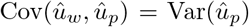 for prediction using the conditional mean, which may be non-linear. Let

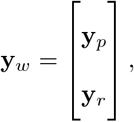

where y_*w*_, y_*p*_, and y_*r*_ indicate a vector of phenotype records in the whole, partial, and validation data set, respectively. It is convenient to first show that 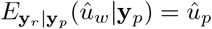:

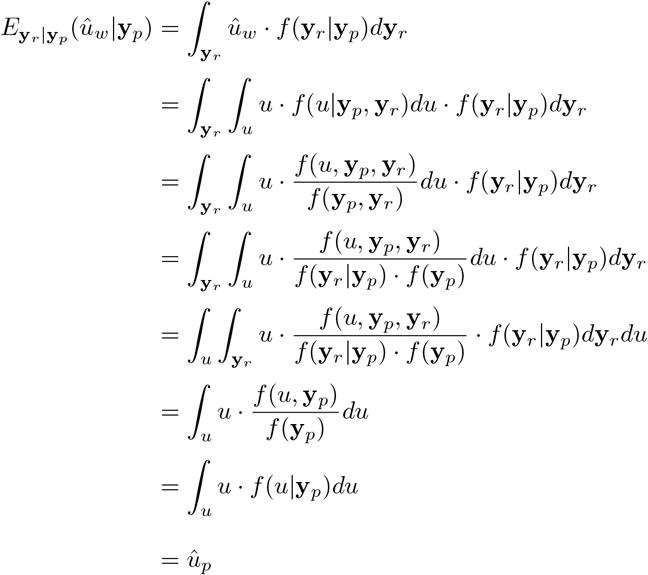

Now, we write the 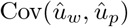 as:

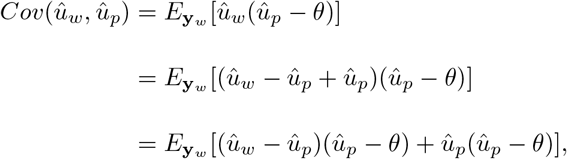

where *θ* is the expected value of 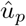. But the first term of this expectation can be shown to be null:

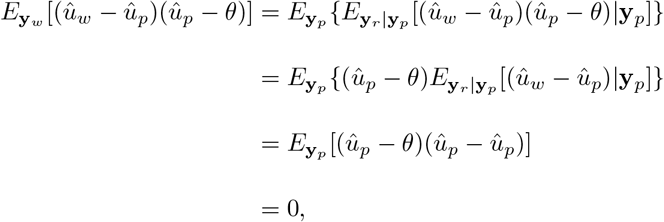

because, as shown previously, 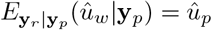. Thus, the 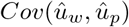 becomes:

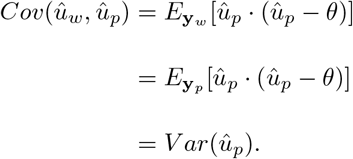

With the proof of 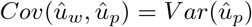, we showed the LR method holds for non-linear models. This proof is similar in principle to that provided by Belay et al. [15], but we recognize that it is not limited to BLUP, as invoked in that study, but is applicable to any method of prediction based on the conditional mean [16], including for non-linear models.

## Data simulation

A longitudinal data set of body weights in pigs was simulated to evaluate the behavior of LR method for non-linear models, both when the true and a wrong model are used for analysis. Body weights of 1500 individuals from 70 to 500 days of age were simulated using a combination of multi-trait QTL effects (30 bi-allelic QTL) and a Gompertz growth model. Following van Milgen et al. [17], the body weight of individual *i* at age *t* (*BW_it_*) was simulated as:

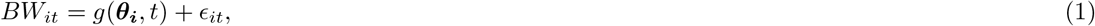

where *θ_i_* = [*Age115*_*i*_ *Shape*_*i*_ *BW* 65_*i*_] refers to three underlying latent variables for pig *i* of age at 115 kg, a shape parameter, and body weight at 65 days, and *ϵ_it_* is the residual. We simulated a heterogeneous residuals to mimic the real growth data for pigs using three different residuals across days 70 to 500 (i.e., 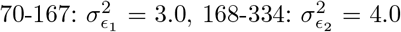, and 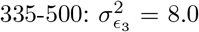). In equation (1), *g*(.) indicates the nonlinear Gompertz function [17]:

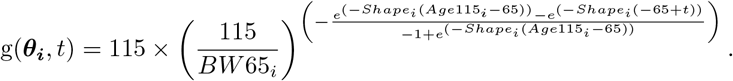

The three underlying latent variables *θ_i_* for individual *i* were considered correlated and modeled with a multivariate QTL effects model.

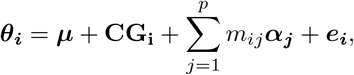

where *μ* is a vector with the intercepts for each latent variable, **CG_i_** is a vector of contemporary group effects, *m_ij_* is the genotype covariate (0, 1, 2) of individual *i* at the *jth* QTL, *α_j_* is a vector of effects for the three latent variables for the *jth* QTL, and *e_i_* is a vector of random environmental effects associated with each latent variable. Based on the results of Yu et al. [18], the variance-covariance matrix used for simulation of random environmental effects was equal to 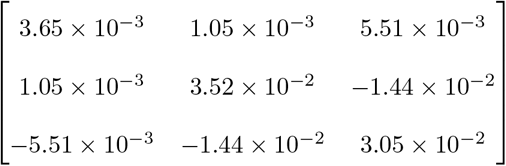. The variance-covariance used to simulate QTL effects for the three latent variables was arbitrarily but without loss of generality derived by dividing the environmental variance-covariance by the number of QTL (i.e., 30).

Using the QTL as markers, the simulated data were analyzed with two Bayesian Hierarchical models: 1) the Gompertz model that was used for simulation, i.e. the true model, and 2) a quadratic growth model, i.e. a wrong model. Variance components that were used to simulate the data were fitted into the true and wrong models for analysis. The prediction performances of these two models were evaluated using the LR method across 20 replicates. All analyses were performed in Julia [19].

## Data analysis models

We analyzed the simulated data using the Bayesian Hierarchical Gompertz growth model (BHGGM) developed by Yu et al. [18], which integrates a Gompertz growth model, i.e. the true model, with a multi-trait marker effects models. Following equation (1), the three underlying latent variables in the Gompertz growth model were assigned the following prior:

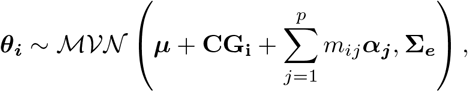

where Σ_e_ is the environmental variance-covariance matrix, which was assumed to have an inverse Wishart prior, 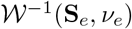. The prior for *ϵ_it_* had a null mean and age specific variances (as described above) to allow fitting heterogeneous residuals. Flat priors were assigned to *μ* and **CG_i_** and the prior for *α_j_* followed 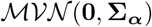, where Σ_*α*_ has an inverse Wishart distribution, 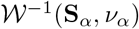.

To fit the Bayesian Hierarchical quadratic growth model (BHQGM), i.e. the wrong model, we introduced a quadratic growth model for the non-linear function *g*(.) in equation (1):

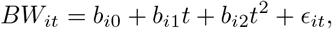

where *b*_*i*0_, *b*_*i*1_, and *b*_*i*2_ refer to three underlying latent variables for individual i and were assigned the same multivariate normal prior as *θ_i_* in BHGGM. The other parameters in BHQGM also used the same priors as in BHGGM.

## Design of the partial and validation data sets

To investigate the behavior of the LR method, three partitioning scenarios (Figure 1) were implemented: 1) by animal: phenotype records for days 70 to 500 of the first 500 individuals comprised the partial set and the phenotypes of the remaining 1000 individuals were assigned into the validation set, 2) by age: phenotypes for days 70 to 300 of all 1500 individuals comprised the partial set and all 1500 individuals and their phenotypes from days 301 to 500 were considered as the validation set, and 3) by animal and age: phenotypes for days 70 to 300 of the first 500 individuals comprised the partial set and phenotypes for days 301 to 500 for the remaining 1000 individuals were assigned to the validation set. The EBV of body weights for individuals in the validation set across days were then predicted based on the partial set of different scenarios.

**Figure 1:**
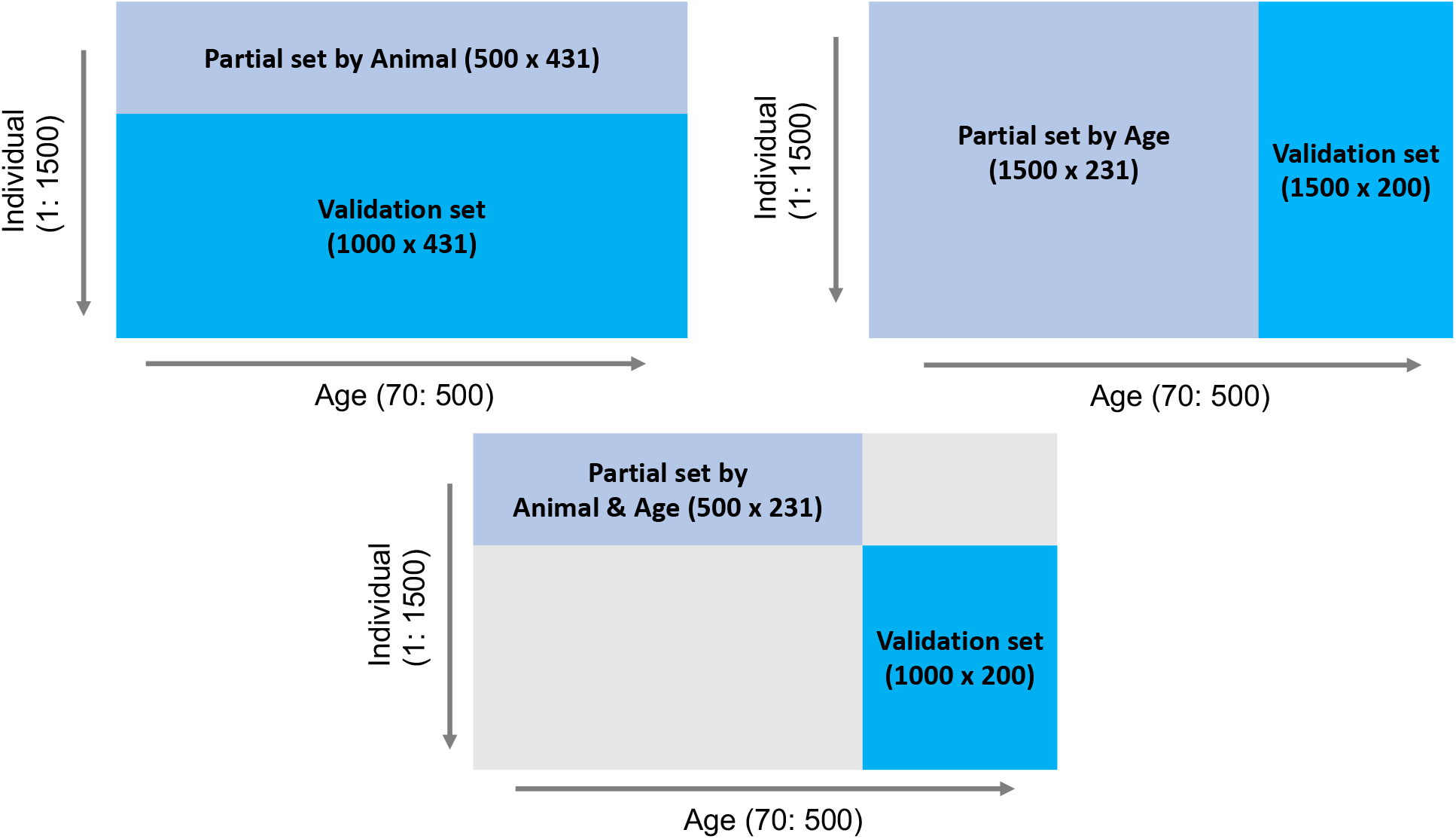
Outline of three data partitioning scenarios to create partial and validation sets for the LR method.

### LR method estimators

Following Legarra and Reverter [11], estimators of bias, inflation, and accuracy were calculated for EBV of body weights of the individuals in the validation set using the LR method, as described below.

1. The LR estimator of population bias is 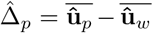, which is the difference between the mean EBV of individuals in the validation set estimated from partial and whole data sets, where a value of 0 indicates no bias.
2. The estimator of inflation is obtained by computing the regression coefficient of 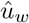 on 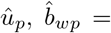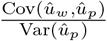, which is an estimator of the regression of true breeding value on 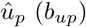. This estimator has been referred to as estimator of dispersion by Legarra and Reverter [11] and inflation by Belay et al. [15], where 1 indicates no inflation. Suppose animals are selected based on EBV to increase the values of a trait. Then if the true inflation is less than 1, the BV of selected candidates is expected to be lower than their EBV, which indicates an upward bias of the EBV of the selected animals. On the other hand, when 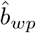 is larger than 1, the BV of selected candidates will be higher than their EBV, which indicates an downward bias of the EBV of the selected animals.
3. The estimator of population accuracy, 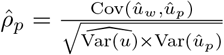, where 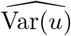 refers to an estimate of the genetic variance of individuals in the validation set. This estimate was obtained by Gibbs sampling as: 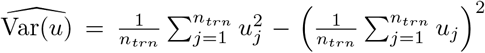, where *u_j_* refers to the sampled breeding value of individual *j* and *n_trn_* is the total number of individuals in the training set.

In addition to these LR estimators of bias, inflation, and accuracy for body weights predictions, we also calculated the “true” estimators of these parameters using the simulated values of *u* in place of 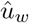 for each day. Note that these “true” estimators can only be computed in a simulation study, and they are used to study the performance of the LR estimators, which can be computed in real data analyses.

For the true and estimated bias statistics, we calculated their means for each day of age across all animals in the validation set. These bias statistics were averaged across days within each replicate to test whether their mean was significantly different from 0 using a t test. Similarly, true and estimated regression coefficient statistics were averaged across days within each replicate to test whether their mean was significantly different from 1 using a t test.

## Results

To better visualize the prediction performances across the fitted models and partitioning scenarios, we randomly picked one individual from the validation set and displayed its simulated data against its predictions in Figure 2. Both simulated body weight phenotypes, true breeding values, and estimated breeding values of the selected individual were displayed. The predicted data included the breeding values estimated from the partial and whole data sets for the three partitioning scenarios (Figure 2).

**Figure 2:**
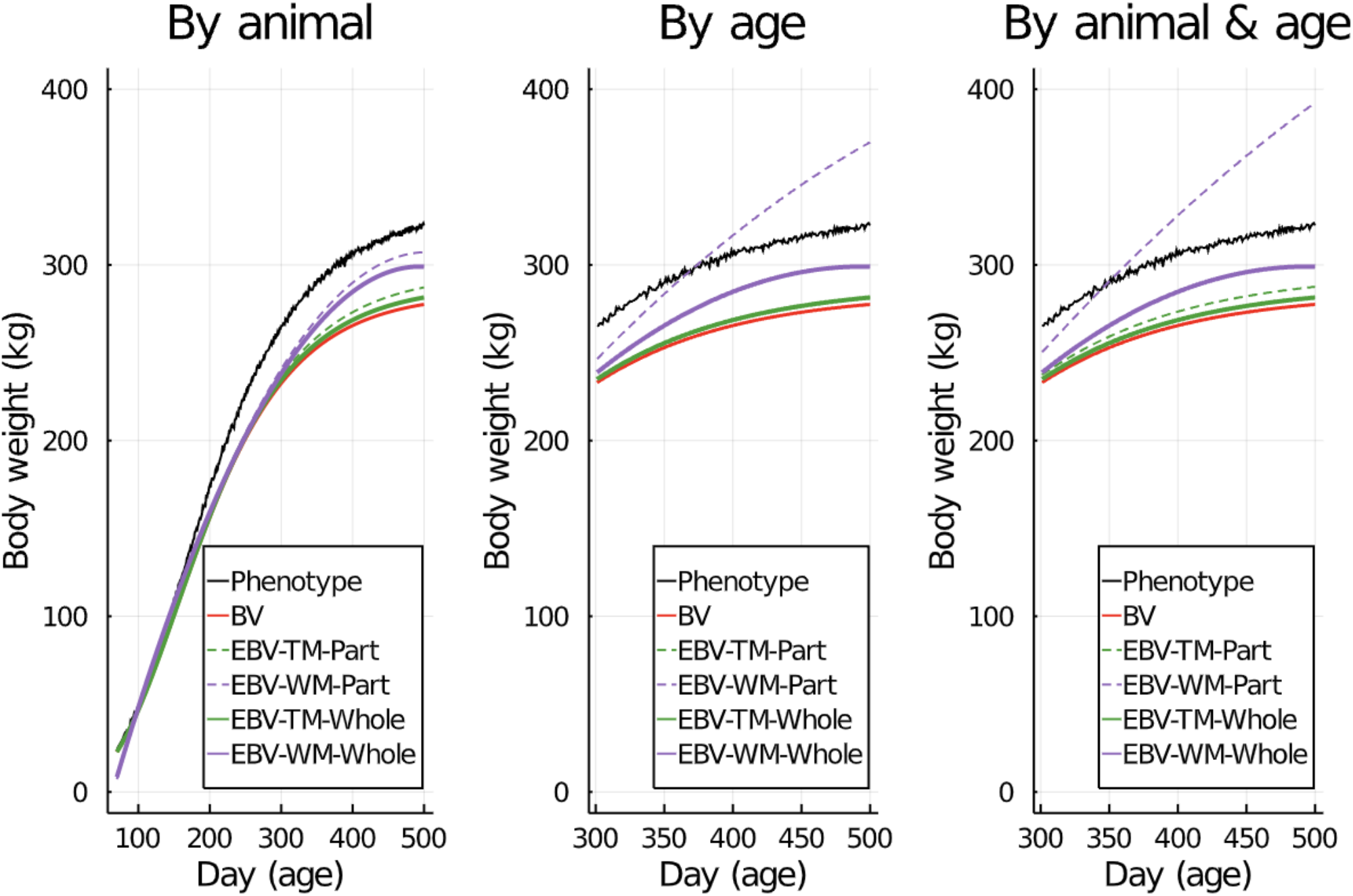
Example of simulated phenotypes, true breeding values (BV), and estimated breeding values (EBV) for body weight by age using the true model (TM) and the wrong model (WM) when the partial data set (Part) was partitioned using different scenarios.

### Population bias

Figure 3 shows the true and LR estimates of prediction bias of EBV for body weight at each day when the data were partitioned by animal. When the true model (BHGGM) was used, both the true and LR estimates of prediction bias were symmetrically distributed around 0 for each day, and their mean was not significantly different from 0 (*P* = 0.84 and *P* = 0.37). In contrast, when the wrong model (BHQGM) was used, the mean of the true estimates of bias was significantly different from 0 (*P* < 0.001), but the LR estimates of bias were symmetrically distributed around 0 for each day and their mean was not significantly different from 0 (*P* = 0.4).

**Figure 3:**
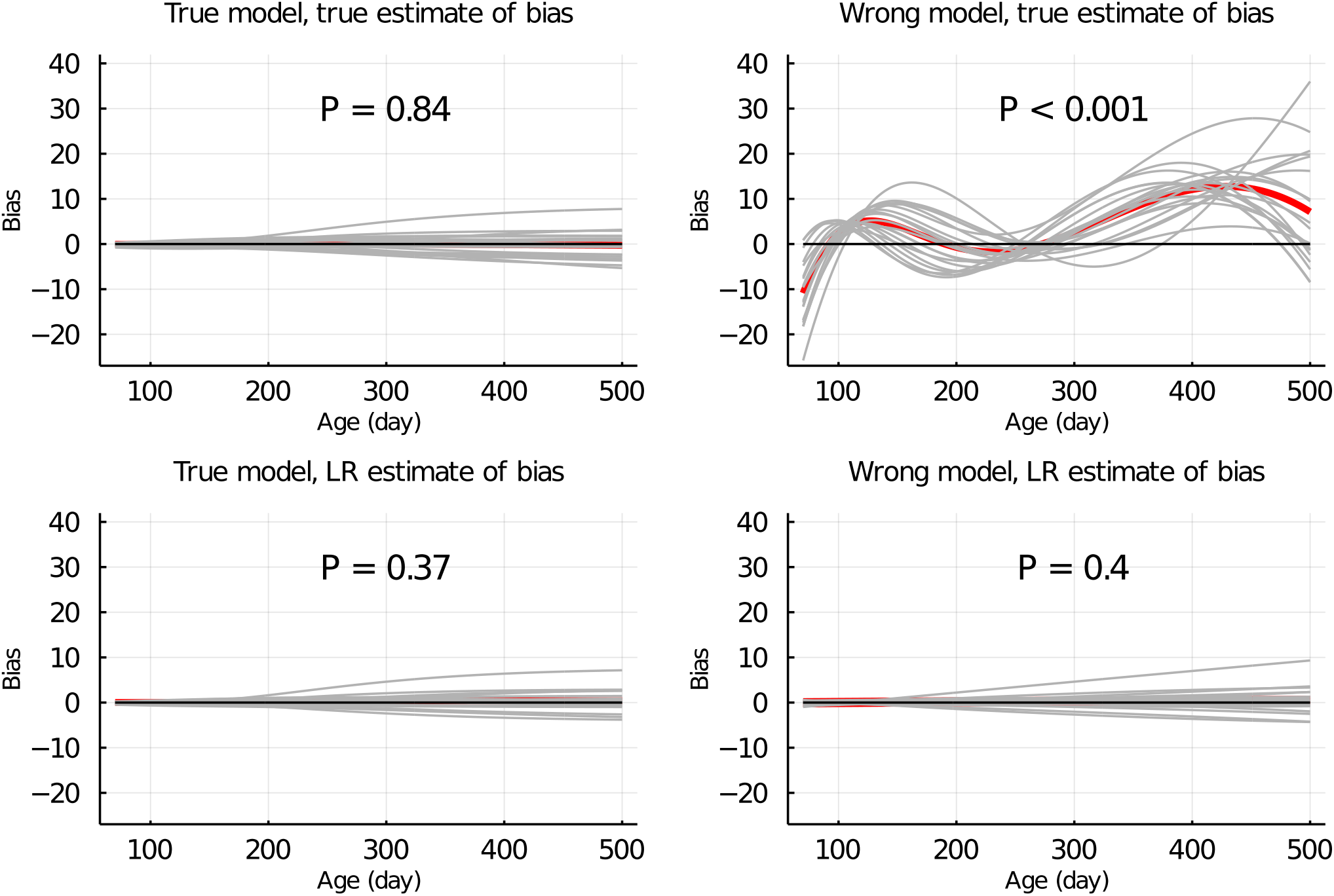
True and LR estimates of bias of EBV of body weights at each day when the true or wrong model was fitted and when partitioning the data by animal. Grey lines are results of 20 simulation replicates, the red line is the mean of 20 replicates, and the black line indicates bias = 0. P refers to significance of tests for the difference between true or LR estimate of bias and 0.

Figure 4 shows the true and LR estimates of prediction bias of EBV for body weights at each day when the data were partitioned by age. When the true model was used, both the true and LR estimates of bias were symmetrically distributed around 0 for each day, and their mean was not significantly different from 0 (*P* = 0.10 and *P* = 0.09). When the wrong model was used, the true and LR estimates of bias were significantly different from 0 (*P* < 0.001 and *P* = 0.002). Results for the partitioning by animal & age were consistent with those in Figure 4 and are shown in Supplemental Figure S1.

**Figure 4:**
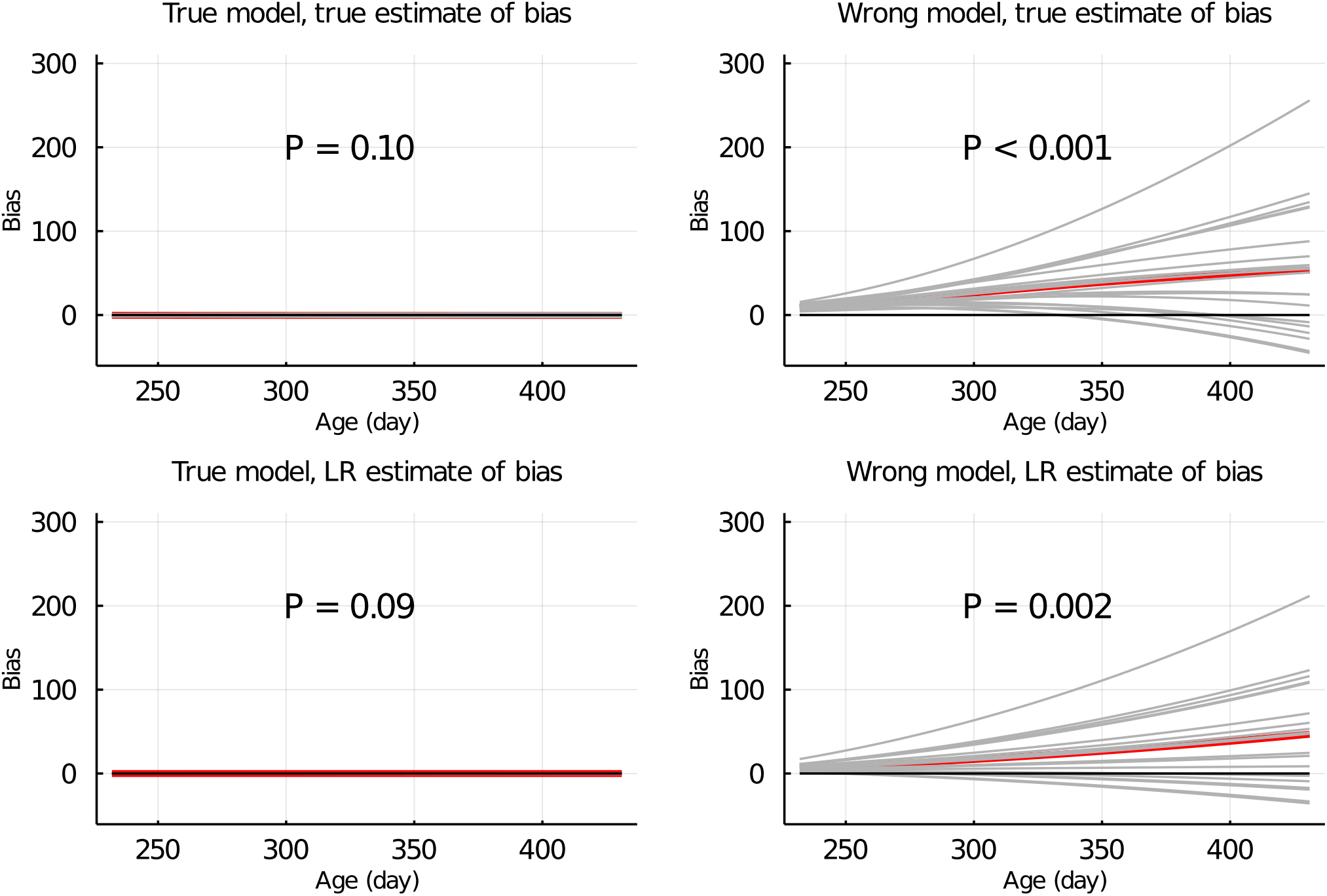
True and LR estimates of bias of EBV of body weights at each day when the true or wrong model was fitted and when partitioning the data by age. Grey lines are results of 20 simulation replicates, the red line is the mean of 20 replicates, and the black line indicates bias = 0. P refers to significance of tests for the difference between true or LR estimate of bias and 0.

Figure 5 shows the true and LR estimates of regression coefficient of EBV for body weights at each day when the data were partitioned by animal. When the true model was used, both the true and estimated regression coefficients were symmetrically distributed around 1 for each day, and their mean was not significantly different from 1 (*P* = 0.75 and *P* = 0.53). When the wrong model was used, the true and LR estimates of regression coefficient were significantly different from 1 (*P* < 0.001). Results for the partitioning by age and by animal & age were consistent with those in Figure 5 and are given in Figure 6 and Supplemental Figure S2, respectively.

**Figure 5:**
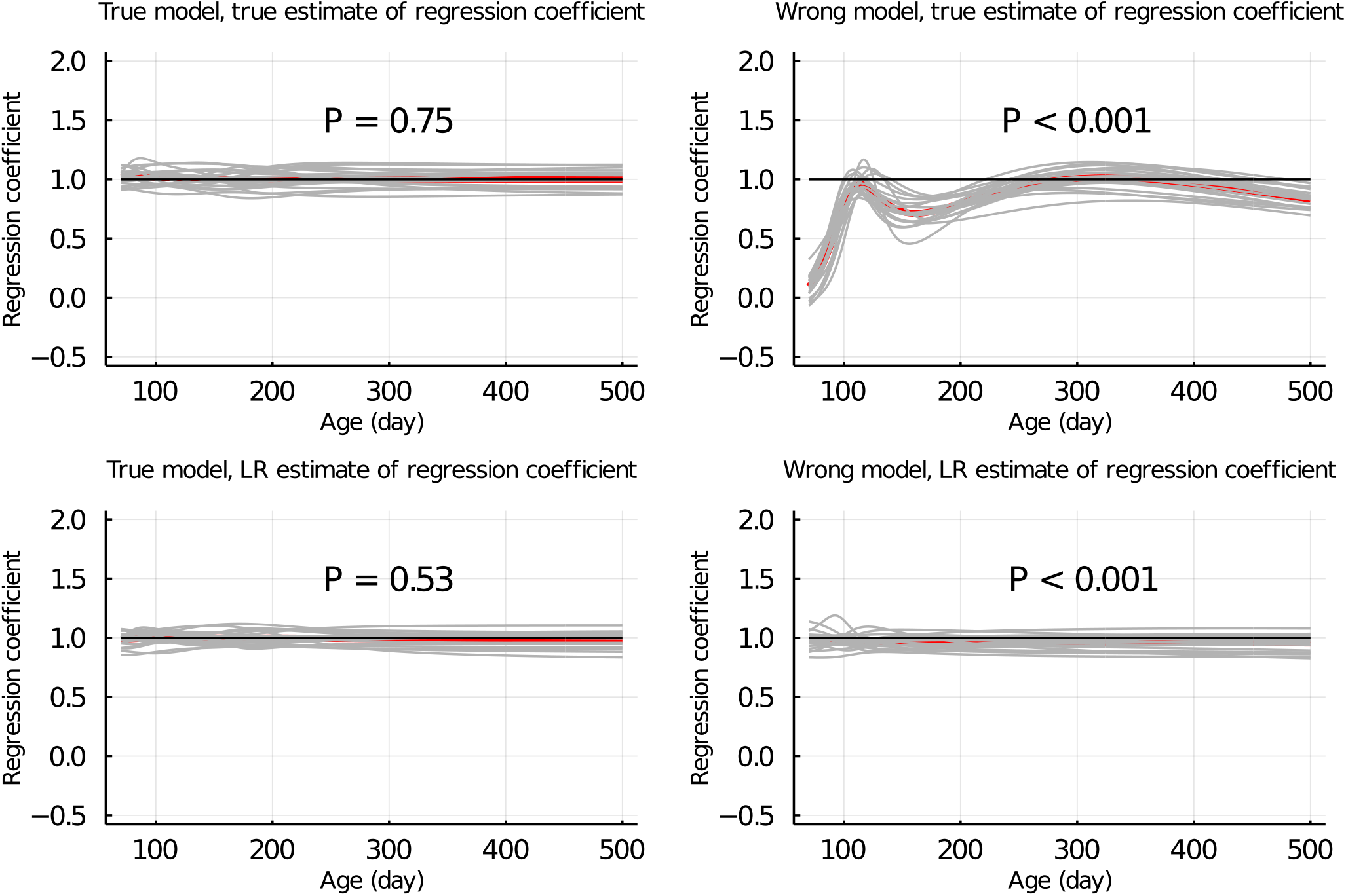
True and LR estimates of regression coefficient of EBV of body weights at each day when the true or wrong model was fitted and when partitioning the data by animal. Grey lines are results of 20 simulation replicates, the red line is the mean of 20 replicates, and the black line indicates regression coefficient = 1. P refers to significance of tests for the difference between true or LR estimate of regression coefficient and 1.

**Figure 6:**
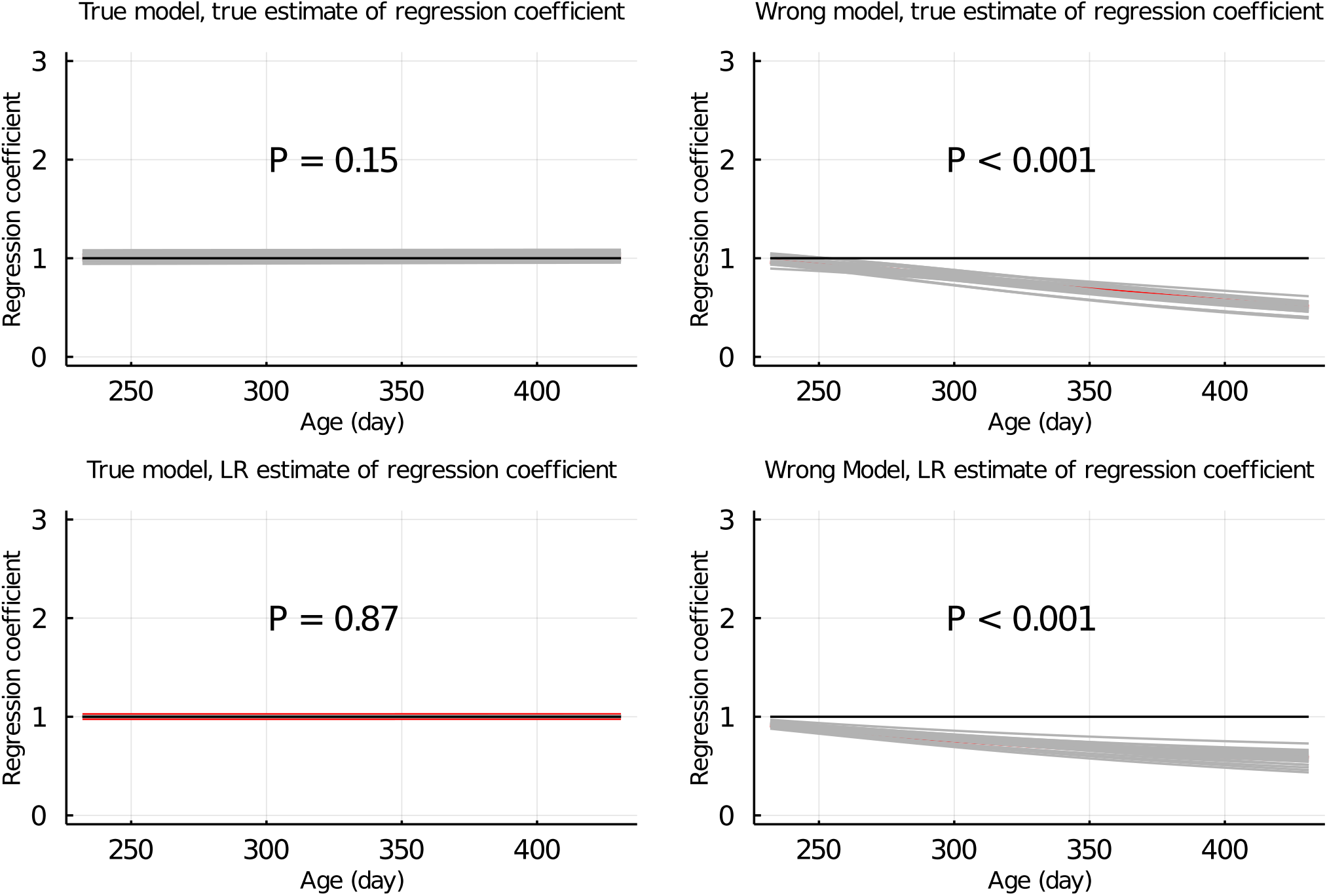
True and LR estimates of regression coefficient of EBV of body weights at each day when the true or wrong model was fitted and when partitioning the data by age. Grey lines are results of 20 simulation replicates, the red line is the mean of 20 replicates, and the black line indicates regression coefficient = 1. P refers to significance of tests for the difference between true or LR estimate of regression coefficient and 1.

### Population accuracy

In Figure 7, the true and LR estimates of prediction accuracy of EBV for body weights at each day when the data were partitioned by animal are presented. The LR estimate of prediction accuracy had a similar pattern as the true estimate of accuracy when using the true model but not when the wrong model was used. When partitioning the data by age, the LR estimate of accuracy showed a similar pattern as the true estimate of accuracy curve regardless of which model was fitted. We also evaluated the difference between 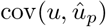 and 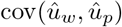 when fitting the true and wrong model for the three data partitioning scenarios (Table 1). There was a non-significant difference (*P* ≥ 0.74) between 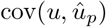 and 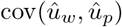 when the true model was fitted, but a significant difference (*P* ≤ 0.004) was observed for each scenario when the wrong model was fitted.

**Figure 7:**
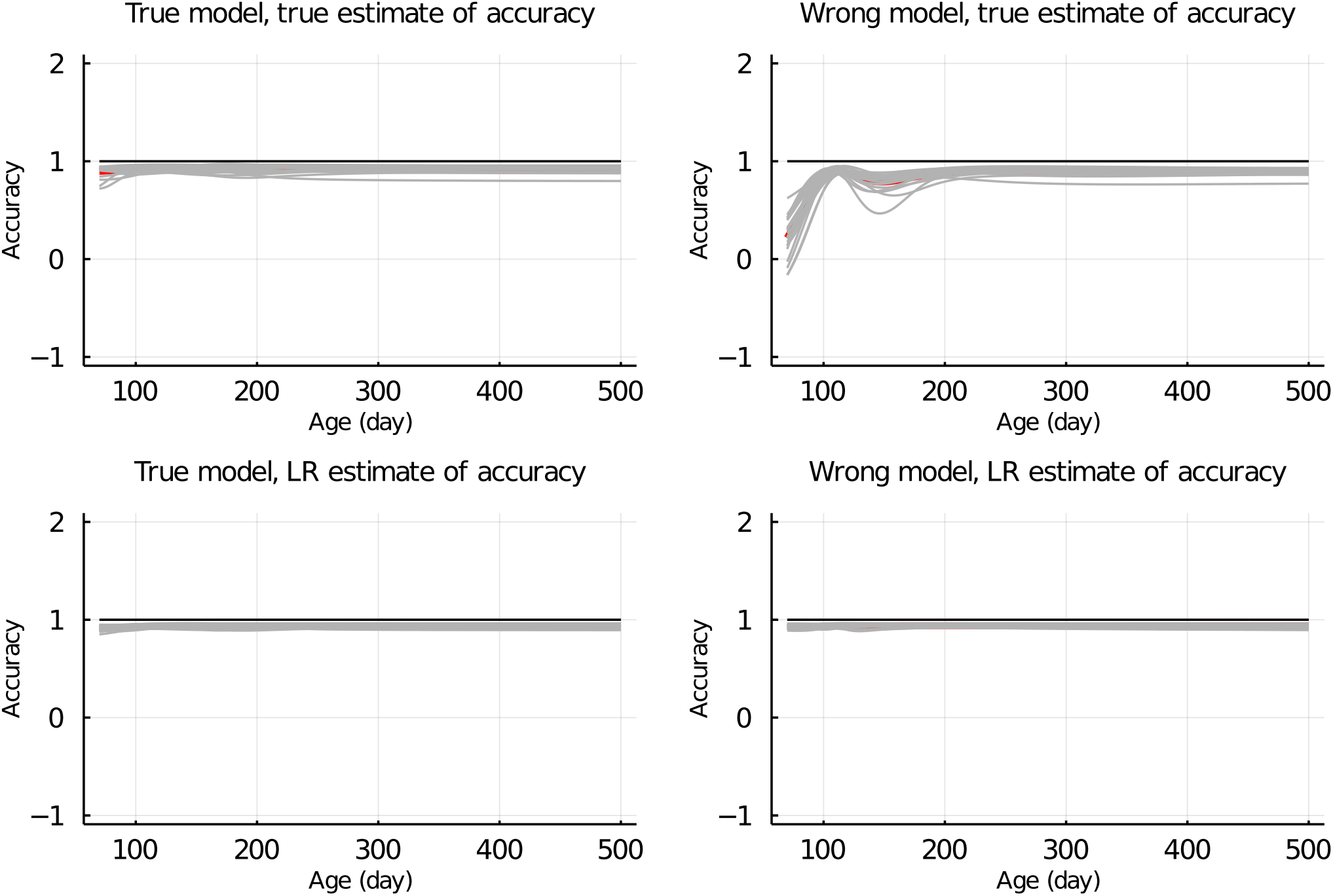
True and LR estimates of accuracy when the true or wrong model was fitted and when partitioning the data by animal. Grey lines are results of 20 simulation replicates, the red line is the mean of 20 replicates, and the black line indicates accuracy = 1.

**Table 1:** Significance (p-values) of tests for the difference between 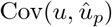 and 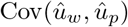 for the three data partitioning scenarios and the two models.

|  | By animal | By age | By animal & age |
| --- | --- | --- | --- |
| True model | 0.74 | 0.85 | 0.79 |
| Wrong model | 0.004 | 0.002 | 0.002 |

## Discussion

Based on the initial idea from Reverter et al. [12], Legarra and Reverter [11] proposed the LR method to quantify the prediction bias and accuracy of EBV at the population level. They proved the validity of LR method for EBV from a linear model using standard BLUP theory and applied the LR method to a real cattle data set [11]. While the LR method has also been applied to EBV from a threshold model [14], a mathematical proof of its validity for a non-linear method of prediction has not been provided. In this study, we presented a mathematical proof for the validity of the LR method for predictions based on the conditional mean [16]. In our proof, we assume the partial data contains a subset of the phenotypes in the whole data. Belay et al. [15] showed the LR method is also applicable to BLUP when the partial data contains a subset of the genotypes in the whole data. The proof presented in the current paper is similar in principle to that provided by Belay et al. [15]. Taken together, these two proofs show that the LR method is applicable to predictions based on conditional mean, regardless of whether the data are partitioned by genotypes or phenotypes and regardless of whether the model is linear or non-linear. Strictly, however, the LR method is only valid if the true model is fitted.

Using simulated longitudinal data, we confirmed our proof and investigated its behavior when a wrong model was used for estimation. Furthermore, we explicitly explored how the strategy for partitioning the data into training and validation sets affect the ability of the LR method to detect bias and estimate accuracy with the model fitted is not the true model, using three different data partitioning strategies. Below, we summarize the implications of the fitted model and data partitioning strategies on the performance of the LR method, thereby providing guidelines for its use to detect bias and estimate accuracy of predictions when a wrong model is fitted.

When the wrong model (BHQGM) was fitted and the data were partitioned by animal, the true estimate of bias was significant, but the LR estimate was not able to identify this bias (Figure 3). Macedo et al. [13] also observed that for a certain misspecification of the model, the LR method was not able to correctly detect and estimate a bias. Figure 5 shows that when the wrong model was used, the true estimate of regression of 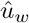 on 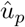 had a significant deviation from 1, and in this case the estimate of the regression coefficient based on the LR method was also significantly different from 1, although differing in magnitude from the true estimate of regression coefficient. This is also consistent with the results observed by Macedo et al. [13]. The pattern of EBV against age were presented in Figure 2 (left column) for a randomly selected individual. When the wrong model was fitted, the EBV from the partial and whole data sets deviated more from the true BV than EBV from the true model did. However, even when the wrong model was used, the EBV from partial and whole data sets were very similar. This explains why the estimate of bias based on the LR method was not significant when the wrong model was used, although there was a true bias. Figure 7 shows that, with the wrong model, the accuracy estimated by the LR method was slightly higher than the true estimate of accuracy, which is consistent with Macedo et al. [13].

When the data were partitioned by age, the LR method was able to correctly detect a bias and inflation when the wrong model was used (Figures 4 and 6). Figure 2 (middle column) shows the EBV of a randomly selected individual when the data were partitioned by animal. When the wrong model was fitted, the EBV estimated from partial and whole data sets both deviated from the true BV but the EBV based on the partial set was quite different from that estimated from the whole set. This illustrates the significant bias that was detected by the LR method for this scenario. Results for the partitioning by animal & age (right column in Figure 2) were similar to those when partitioning by age. As in Macedo et al. [13], even with the use of a wrong model, the accuracy estimated by the LR method was quite close to the true accuracy (Figure 8).

**Figure 8:**
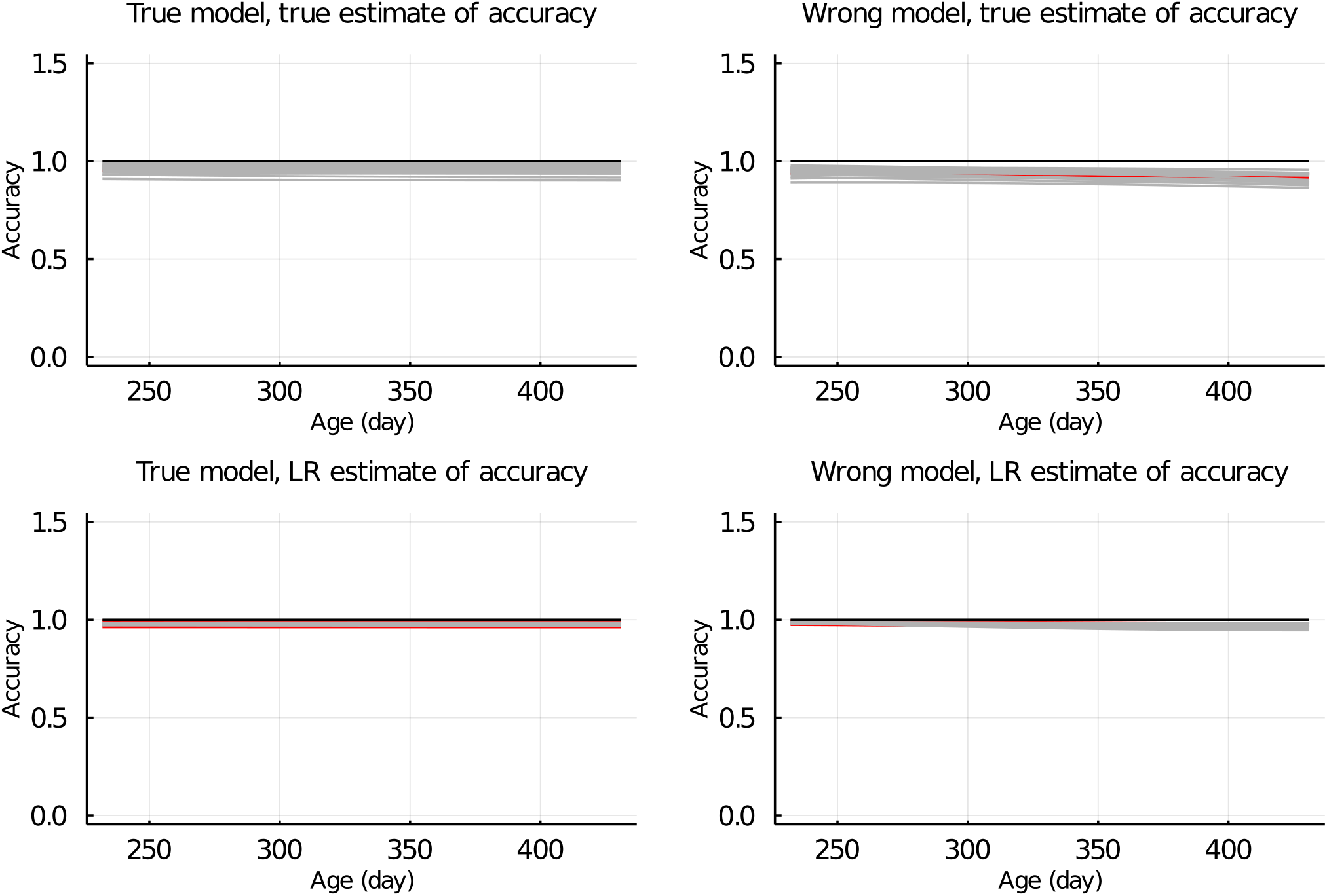
True and LR estimates of accuracy when the true or wrong model was fitted and when partitioning the data by age. Grey lines are results of 20 simulation replicates, the red line is the mean of 20 replicates, and the black line indicates accuracy = 1.

The inconsistent bias estimates obtained with the LR method for different data partitioning strategies suggests that the LR method captures different aspects of the model for different data partitions. When the data were partitioned by animal, both the partial and whole data sets included phenotypes over the range from 70 to 500 days. Thus the fit of the growth curve from the partial and whole data sets were similar, even for the wrong model, although the fit might deviate from that using the true model. Fitting the wrong growth model is only incorrect in the relationship between age and body weight within individual but correctly models relationships between relatives. Thus to appropriately test the predictive ability of the growth model using the LR method, we needed to predict the body weights of animals that are outside the observed age range for animals in the training set. When the data are partitioned by age, the partial data set has only body weights measured at ages up to 300 days, whereas the validation data set has body weights measured at ages up to 500 days. Thus when we predict the body weights of the individuals in the validation set based on the fit of the growth model from the partial data set, we are testing the predictive ability of the growth model. In this case, the LR method was able to correctly detect a bias when using the wrong model (bottom right plots in Figure 4 and Figure 6). In real data analyses, repeated k-fold LR can be used to test the significance of bias. Or if Bayesian method is employed for the LR analysis, the posterior probability of bias can be computed from a single partitioning of the data.

In general, to properly test the predictive ability of a model with the LR method, we need to use the model to predict the performance of individuals that have values for the relevant predictor variables or combination of predictor variables that were not present in the training data. In our simulated data, the predictor variables included the marker genotypes, as well as age. Let’s define the predicted performance of individual *i* as 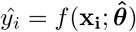, where *f*(.) is the linear or non-linear function used for prediction, **x_i_** is a vector of predictor variables for individual *i*, and 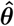 is estimates of model parameters. Below we will use genomic prediction by ridge regression BLUP (RR-BLUP) as an example for illustration. To evaluate the predictive ability of RR-BLUP, the data are partitioned into training and validation sets. The training set is used to fit the predictive model *f*(.) and to estimate the model parameters *θ* (i.e., marker effects). By plugging the marker effect estimates 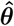 and observed marker genotypes x into *f*(.), the performance of individuals in the validation set can be predicted. The predictive ability of the model can then be quantified by comparing the predicted and observed performances of individuals in the validation set. In RR-BLUP, the relevant predictor variables are the marker genotypes. Thus the same records cannot be used in both the training and validation sets. In our example, the LR method was used to determine whether it could detect a bias when a wrong model was used for analysis of longitudinal body weight data. When predicting longitudinal body weights, the relevant predictor variable is not the genotype but the age of the animal. When the data were partitioned by animal, the training (partial) and validation sets included phenotypes for animals with age ranging from days 70 to 500, the same age range as training data was used for the validation data and, therefore, the LR method failed. However, when the data were partitioned by age, the model was trained using phenotypes with ages ranging from days 70 to 300 and it was tested by predicting body weights for animals with age ranging from days 301 to 500. In this case, the LR method was able to detect a bias when using the wrong model. This was even true when the same genotypes were used in both the training and validation sets, because to check if the model used for predicting longitudinal body weights is correct, the relevant predictor variable is age.

## Conclusions

In the present study, we provide a mathematical proof for the validity of applying the LR method to predictions based on the conditional mean, regardless of whether it is a linear or non-linear function of data. Using simulated data, we observed that the LR method was able to detect bias in predictions when an incorrect non-linear model was fitted. However, when a wrong model is fitted, testing the predictive ability of the model using the LR method is only valid if the validation set includes values of relevant predictor variables that are not present in the training set. To our knowledge, this marks the first study that provides a mathematical proof of the validity of using LR method to a non-linear method of prediction, and we provide guidelines on how to partition data such that the LR method can detect bias and estimate accuracy of predictions when the model fitted is incorrect.

## Authors’ contributions

HY, JCMD, and RLF conceived the research idea. HY and RLF derived a mathematical proof for the validity of the LR method for predictions based on conditional mean. HY performed the data analyses and drafted the manuscript. JCMD and RLF edited the manuscript. All authors read and approved the final manuscript.

## Ethics declarations

### Ethics approval and consent to participate

Not applicable.

### Consent for publication

Not applicable.

---

**Figure S1:**
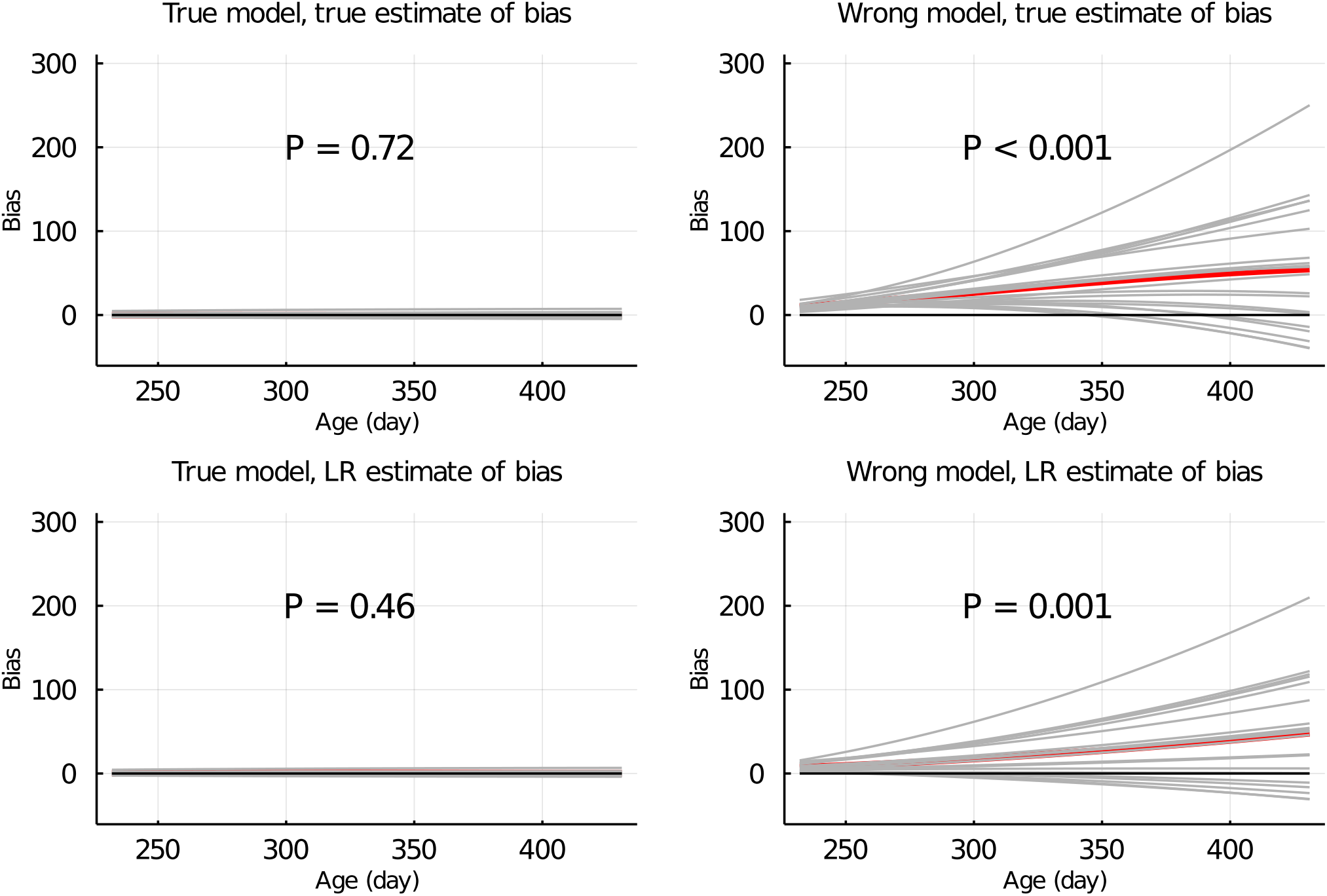
True and LR estimates of bias of EBV of body weights at each day when the true or wrong model was fitted and when partitioning the data by animal & age. Grey lines are results of 20 simulation replicates, the red line is the mean of 20 replicates, and the black line indicates bias = 0. P refers to significance of tests for the difference between true or LR estimate of bias and 0.

**Figure S2:**
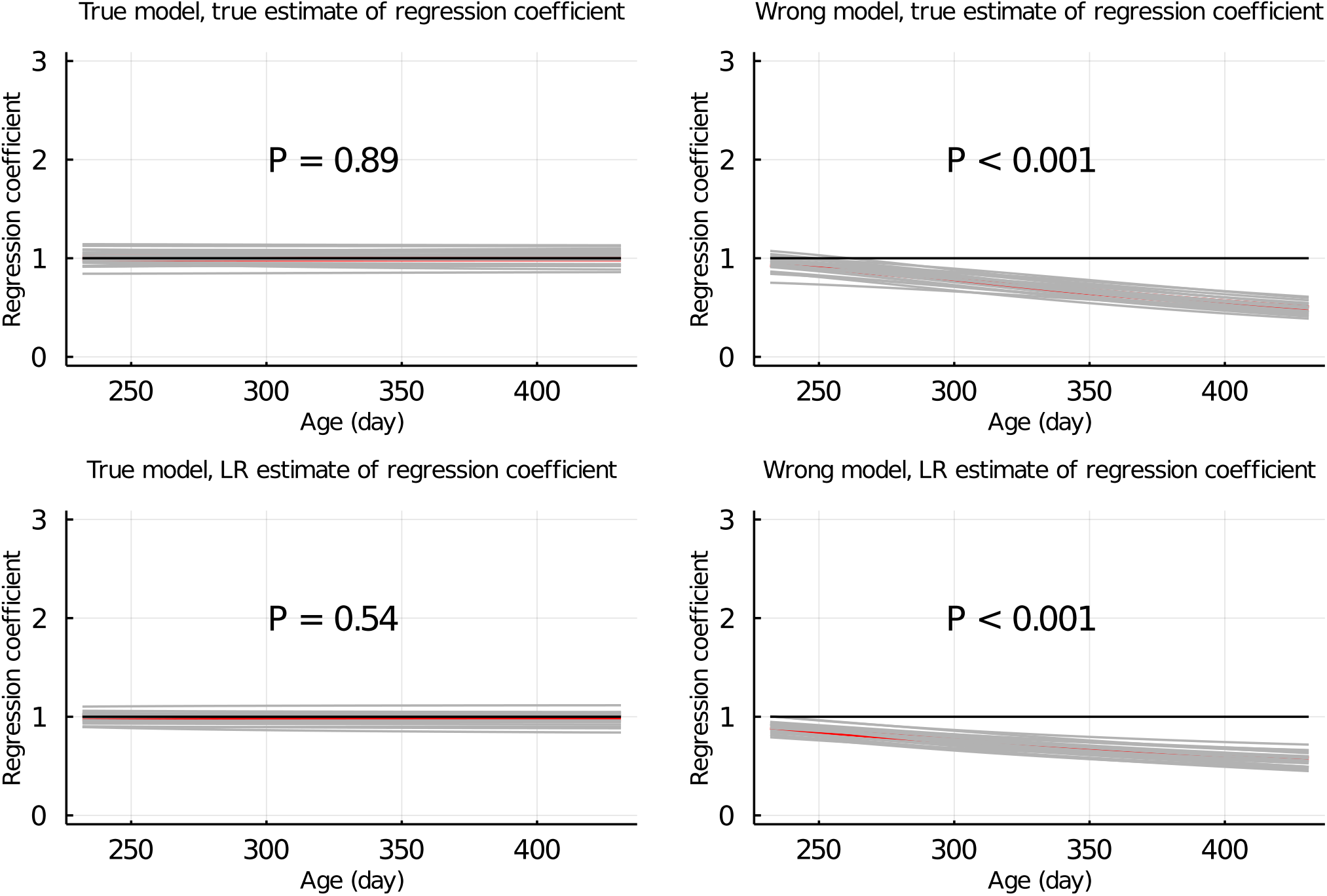
True and LR estimates of regression coefficient of EBV of body weights at each day when the true or wrong model was fitted and when partitioning the data by animal & age. Grey lines are results of 20 simulation replicates, the red line is the mean of 20 replicates, and the black line indicates regression coefficient = 1. P refers to significance of tests for the difference between true or LR estimate of regression coefficient and 1.

**Figure S3:**
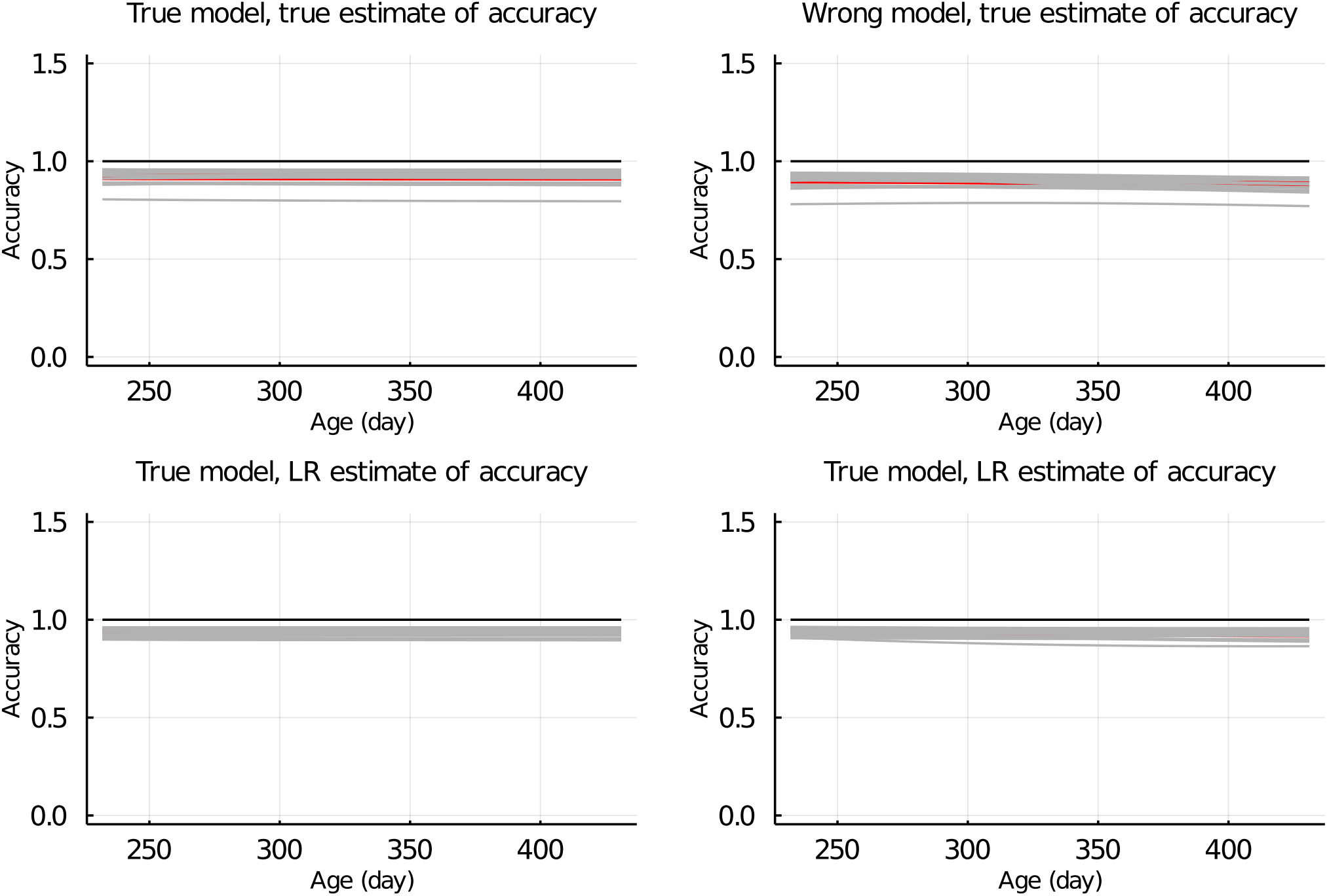
True and LR estimates of accuracy of EBV of body weights at each day when the true or wrong model was fitted and when partitioning the data by animal & age. Grey lines are results of 20 simulation replicates, the red line is the mean of 20 replicates, and the black line indicates accuracy = 1.

